# Proteasomal regulation of ASK family kinases dictates cell fate under hyperosmotic stress

**DOI:** 10.1101/2023.05.23.541899

**Authors:** Xiangyu Zhou, Kengo Watanabe, Kazuhiro Morishita, Jun Hamazaki, Shigeo Murata, Isao Naguro, Hidenori Ichijo

## Abstract

The proteasome has an essential role in proteostasis maintenance and is critical for cell survival under proteotoxic conditions including hyperosmotic stress. However, it is unknown how proteasome activity is linked to cell survival/death under hyperosmotic stress. We have previously reported that apoptosis signal-regulating kinase 3 (ASK3) contributes to cell survival through its inactivation under hyperosmotic stress. 19S regulatory particle subunits of the proteasome were enriched in the ASK3 inactivator candidates identified through our genome-wide small interfering RNA (siRNA) screen. In this study, we demonstrate that the proteasome regulates ASK3 inactivation through its proteolytic activity. Intriguingly, the proteasome inactivates ASK3 via degradation of not ASK3 per se but another ASK family member ASK1 which activates ASK3 in a kinase activity-dependent manner. Furthermore, the elevated ASK1 level under proteasome inhibition sensitizes cells to hyperosmotic stress. These findings suggest that ASK family maintenance links proteasome capacity to cell fate bifurcation under hyperosmotic stress.

## Introduction

Cells are separated from the outside environment by the plasma membrane. Since it is impermeable to diverse biomolecules and has a relatively high permeability to water ^1^, osmotic balance between intracellular and extracellular space is critical for water flux across the plasma membrane. When extracellular osmolality is higher than intracellular osmolality, water flows from intracellular to extracellular space, resulting in cell shrinkage. This condition, hereinafter referred to as hyperosmotic stress, has deleterious effects on the structure and function of biomolecules, organelles, and cytoskeleton ^2^. Upon hyperosmotic stress, cells rapidly induce a volume recovery system known as regulatory volume increase (RVI), but excessive and persistent cell shrinkage leads to apoptosis ^3,4^.

Cell shrinkage under hyperosmotic stress leads to increases in intracellular ionic strength and macromolecular crowding ^3^, which can be a crisis for protein homeostasis (i.e., proteostasis). In vitro studies have suggested that increased ionic strength changes the stability of protein structure and decreases protein activity ^5–7^ and that increased macromolecular crowding accelerates the aggregation of partially unfolded proteins ^8–10^. Consistently, hyperosmotic stress induces protein misfolding and aggregation in intact cells ^11,12^. These reports suggest that hyperosmotic stress increases the burden on proteostasis machineries that degrade abnormal intracellular proteins, such as the proteasome and autophagy. In addition, disruption of the proteasome or autophagy decreases survival of *C. elegans* and mammalian cells under hyperosmotic stress ^13–15^, suggesting that the integrity of proteostasis machineries is a key factor for the survival of cells faced with proteotoxicity under hyperosmotic stress. However, molecular mechanisms which link the overloaded condition of cellular proteostasis machineries to cell survival under hyperosmotic stress remain obscure.

Previously, we found that apoptosis signal-regulating kinase 3 (ASK3; also known as mitogen-activated protein kinase kinase kinase 15 [MAP3K15]) is rapidly inactivated upon hyperosmotic stress ^16^ and that its inactivation is important for RVI and cell survival under hyperosmotic stress ^17^. By leveraging a genome-wide small interfering RNA (siRNA) screen, we have reported upstream regulators of ASK3 inactivation ^17,18^. At the same time, we noticed that many proteasome-constituent genes were retained as the ASK3 inactivator candidates by the screen.

In the present study, we report that the proteasome is a regulator of ASK3 inactivation under hyperosmotic stress. We demonstrate that proteasome activity downregulates the protein level of another ASK family member, ASK1 ^19^, and thereby suppresses the inhibitory effect of ASK1 on ASK3 inactivation. Furthermore, ASK1 mediates the acceleration of hyperosmotic stress-induced apoptosis under proteasome inhibition. Our results suggest that proteasomal regulation of ASK1 protein levels controls ASK3 signaling and cell death under hyperosmotic stress, pointing out the role of ASK family proteins as linking molecules between proteostasis capacity and cell fate to hyperosmotic stress.

## Results

### The Proteasome Negatively Regulates ASK3 Activity under Hyperosmotic Stress

In our previous study, a high-content genome-wide siRNA screen was conducted to comprehensively identify the negative regulators of ASK3 activity under hyperosmotic stress and 63 positive genes were obtained from the screen (^17^; Figures S1A and S1B). We noticed that many proteasome subunit genes were included in the positive genes (Figures 1A and 1B). We then performed functional enrichment analysis on the positive genes and found that genes encoding the 19S regulatory particle (RP; PA700 in Figure 1B) of the proteasome, were significantly enriched in the positive genes (Figures 1B, 1C, and S1C). To validate the screen results, we examined the knockdown effects of the positive genes encoding 19S RP subunit on ASK3 activity, using a phospho-specific antibody that recognizes Thr808 of ASK3, whose phosphorylation is critical for its kinase activity ^16^. Depletion of proteasome 26S subunit, ATPase 2 (PSMC2) upregulated the activity of exogenously expressed ASK3 under hyperosmotic stress, detected by immunofluorescence (Figures 1D). The upregulated ASK3 activity under hyperosmotic stress by PSMC2 depletion was also confirmed by immunoblotting for both exogenously expressed ASK3 (Figure 1E; lanes 2, 4, and 6) and endogenous ASK (Figure 1F; lanes 2, 4, and 6). Note that the phospho-specific antibody for ASK3 also recognizes phosphorylated form of other ASK family proteins^20^ and reflects overall ASK activity derived from ASK1 and ASK3 due to the similar and relatively high molecular weight (ASK1: 155 kDa, ASK3: 147 kDa) particularly in endogenous-level immunoblotting evaluations (e.g., Figure 1F). Depletion of another 19S RP subunit, PSMC3, also resulted in increased endogenous ASK activity in hyperosmotic stress (Figure S1D). These results suggest that the proteasome suppresses ASK3 activity under hyperosmotic stress.

**Figure 1.**
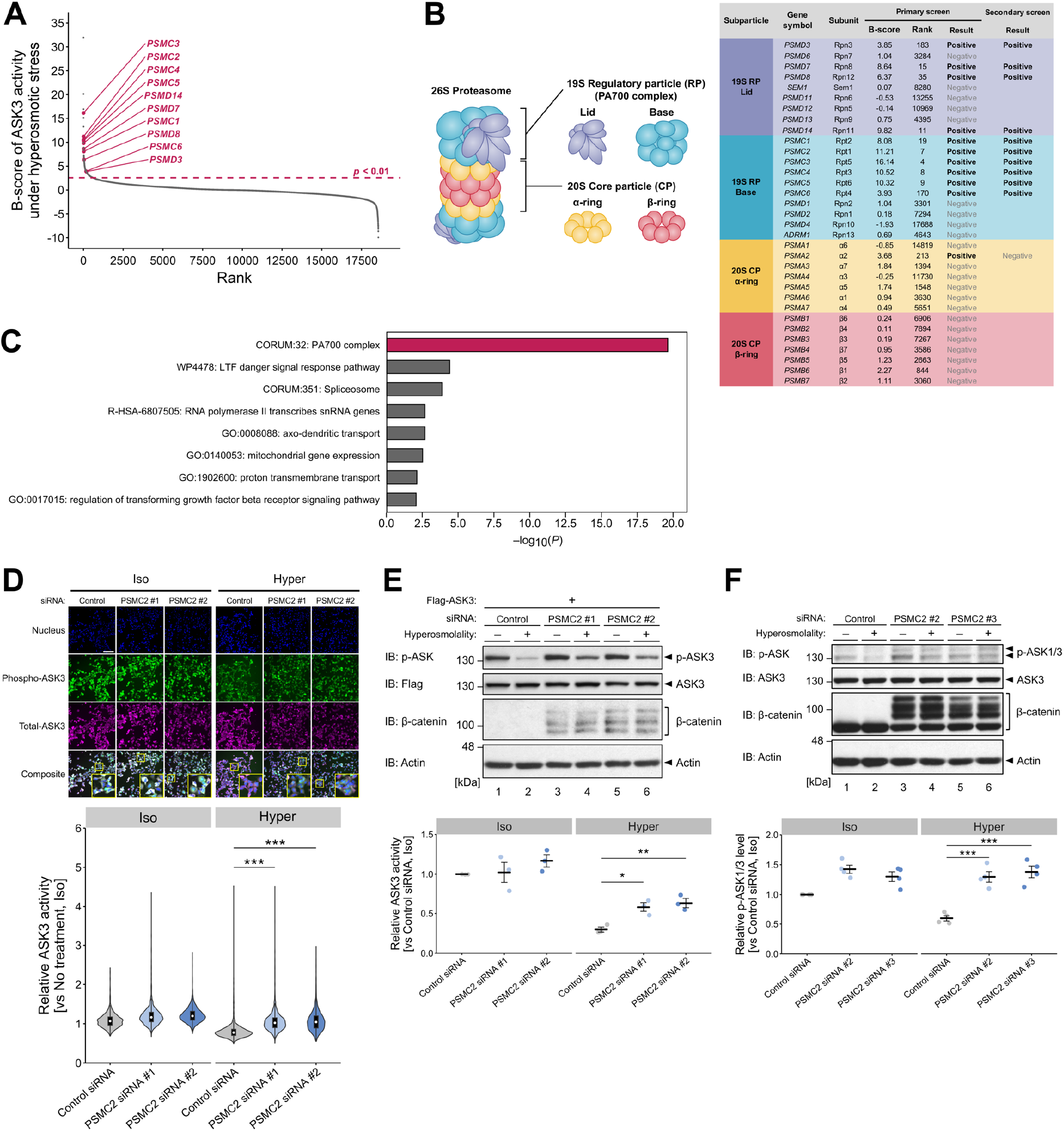
The Proteasome Is a Regulator of ASK3 Inactivation under Hyperosmotic Stress. (A) A rank-ordered plot of *B*-scores for all genes subjected to the primary screen ^17^. Each gene is represented as each point on the plot. A gene with a higher *B*-score corresponds to a stronger ASK3 inactivator candidate. The 19S proteasome subunit genes that were positive on the screen are highlighted in magenta. (B) Scores, ranks, and results of proteasome subunit genes in the genome-wide siRNA screen. (C) Gene Ontology terms enriched in the positive genes of the secondary screen. Terms satisfying the default criteria of Metascape analysis ^38^ and their nominal *p*-values are presented. 19S RP (PA700 complex) is highlighted in magenta. (D) Effects of PSMC2 depletion on ASK3 activity in tetracycline-inducible Flag-ASK3-stably expressing cells. The top panel shows immunofluorescence images of nuclei, phospho-ASK3, total-ASK3, and their composite with enlarged image of squared region. The bottom panel shows violin plots of ASK3 activity in each cell. ASK3 activity was defined as the relative fluorescence intensity of phospho-ASK3 to total ASK3. Control siRNA: *n* = 1,065 (iso), 1,493 (hyper); PSMC2 siRNA #1: *n* = 1,943 (iso), 2,234 (hyper); PSMC2 siRNA #2: *n* = 2,000 (iso), 2,293 (hyper). Representative data from three independent experiments. The white scale bar represents 200 µm. (E, F) Effects of PSMC2 depletion on the activity of exogenously expressed ASK3 (E) or endogenous ASK3 (F) under hyperosmotic stress. Accumulation of ubiquitinated β-catenin is shown as an efficiency indicator of proteasome dysfunction due to PSMC2 depletion. The bottom graphs depict the quantification of western blots. Individual values and the mean ± SEM are presented as points and bars, respectively. *n* = 3 (E) and 4 (F). (D–F) Iso, 300 mOsm; Hyper, 400 mOsm; 10 min. IB, immunoblotting. \**p* < 0.05, \*\**p* < 0.01, \*\*\**p* < 0.001. See also Figure S1.

### Proteasome Activity Is Required for ASK3 Inactivation under Hyperosmotic Stress

The proteasome is composed of two subcomplexes: the catalytic 20S core particle (CP) and the 19S RP. In the process of ubiquitin-dependent degradation, the 19S RP recognizes, unfolds, deubiquitinates, and translocates the ubiquitinated substrates to the 20S CP where proteolysis takes place ^21^. Interestingly, all proteasome-related genes included in the positive genes were those encoding 19S RP subunits (Figure 1B). We then investigated if the catalytic activity of the proteasome is required for ASK3 inactivation under hyperosmotic stress. Treatment of MG132 and bortezomib, proteasome inhibitors that block catalytic activities of 20S CP subunits^22^, caused significant upregulation of ASK3 activity under hyperosmotic stress (Figures 2A and S2A). Treatment of the proteasome inhibitors upregulated endogenous ASK activity under hyperosmotic stress (Figure 2B). Moreover, siRNA-utilized depletion of a 20S CP subunit PSMA1 also increased ASK activity (Figure S2B), indicating that 20S CP genes were the false negatives in our screen, presumably because the depletion of 20S CP subunit was severely toxic for cells and only cells with low knockdown efficiency survived. These results suggest that the proteolytic activity of the proteasome is required for hyperosmotic stress-induced ASK3 inactivation.

**Figure 2.**
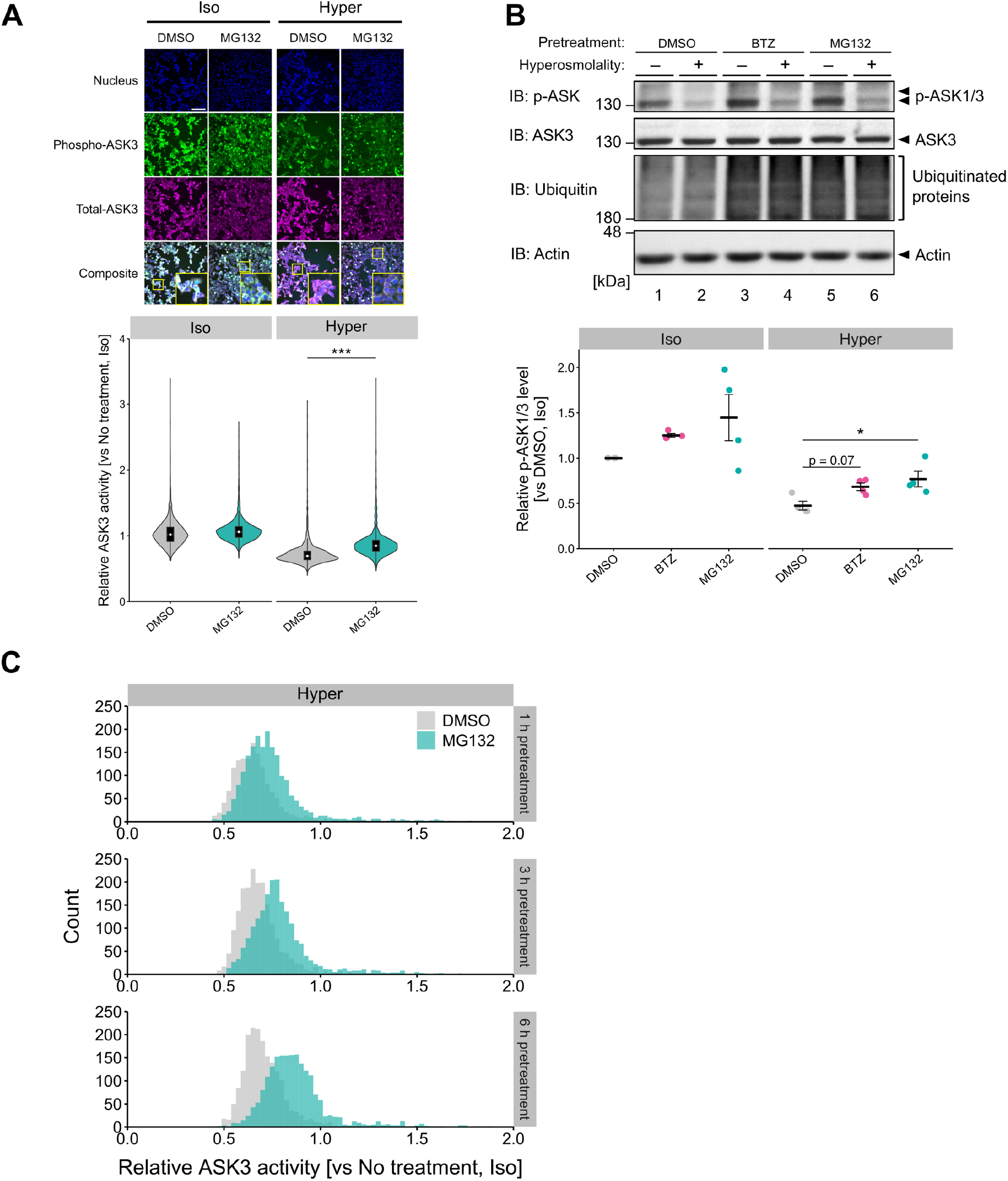
Proteasome Activity Is Required for ASK3 Inactivation under Hyperosmotic Stress. (A) Effects of MG132 on ASK3 activity in tetracycline-inducible Flag-ASK3-stably expressing cells. Cells were pretreated with 10 µM MG132 for 6 h before osmotic stimuli. ASK3 activity was measured as in Figure 1D. The top panel shows immunofluorescence images of nuclei, phospho-ASK3, total-ASK3, and their composite with enlarged images of squared region. The bottom panel shows violin plots of ASK3 activity in each cell. DMSO: *n* = 1,876 (iso), 1,900 (hyper); MG132: *n* = 2,280 (iso), 1,991 (hyper). Representative data from four independent experiments. The white scale bar represents 200 µm. (B) Effects of proteasome inhibitors on endogenous ASK activity. Cells were pretreated with 100 nM bortezomib or 10 µM MG132 for 4 h before osmotic stimuli. Proteasome inhibition efficiency was confirmed by the accumulation of ubiquitinated proteins. The right graph depicts the quantification of western blots. Individual values and the mean ± SEM are presented as points and bars, respectively. *n* = 4. (C) Time-dependency of the MG132 effects on ASK3 activity under hyperosmotic stress. Tetracycline-inducible Flag-ASK3-stably expressing cells were pretreated with 10 µM MG132 for 1/3/6 h before osmotic stimuli. ASK3 activity was measured as in Figure 1D. Histograms of ASK3 activity in each cell are shown. Outlier cells with relative ASK3 activity greater than 2 are not shown. DMSO: *n* = 1,451 (1 h), 1,910 (3 h), 1,900 (6 h); MG132: *n* = 1,964 (1 h), 1,970 (3 h), 1,991 (6 h). Representative data from two independent experiments. (A–C) Iso, 300 mOsm; Hyper, 400 mOsm; 10 min. BTZ, bortezomib; IB, immunoblotting. \**p* < 0.05, \*\*\**p* < 0.001. See also Figure S2.

The MG132 treatment-induced increase in ASK3 activity under hyperosmotic stress (Figure 2A) became greater as the duration of MG132 pretreatment increased from 1 h to 6 h (Figure 2C). At the same time, the proteasome activity decreased to about 10% in 1 h after inhibitor treatment and remained almost constant until 6 h (Figure S2C). These results implied that the time-dependent effect of proteasome inhibitor on ASK3 activity is not due to a time-dependent decrease in proteasome activity, but rather to a time-dependent accumulation of a certain proteasome substrate. Collectively, these results suggest that degradation of proteasome substrate is required for ASK3 inactivation under hyperosmotic stress.

### Increased ASK1 Protein Level Causes a Defect in ASK3 Inactivation under Proteasome Inhibition

We next searched for the proteasome substrate whose degradation is required for ASK3 inactivation. Since the inhibition of protein synthesis by cycloheximide did not decrease the intracellular protein level of ASK3 under hyperosmotic stress within 1 h (Figure S3A), it is unlikely that the hyperosmotic stress-induced ASK3 inactivation stemmed from the rapid proteasomal degradation of the phosphorylated ASK3 (i.e., active form) followed by its replacement with the newly synthesized ASK3 (i.e., inactivated form). In contrast to ASK3, we noticed that the protein level of ASK1 was increased under hyperosmotic stress when the proteasome subunit PSMC3 was depleted (Figure 3A) and when the proteasome activity was inhibited by the bortezomib treatment (Figure 3B). ASK1 mRNA levels were not increased during the bortezomib treatments (Figure S3B), implying that the increased ASK1 protein level under proteasome inhibitions is not due to the transcriptional regulation of ASK1.

**Figure 3.**
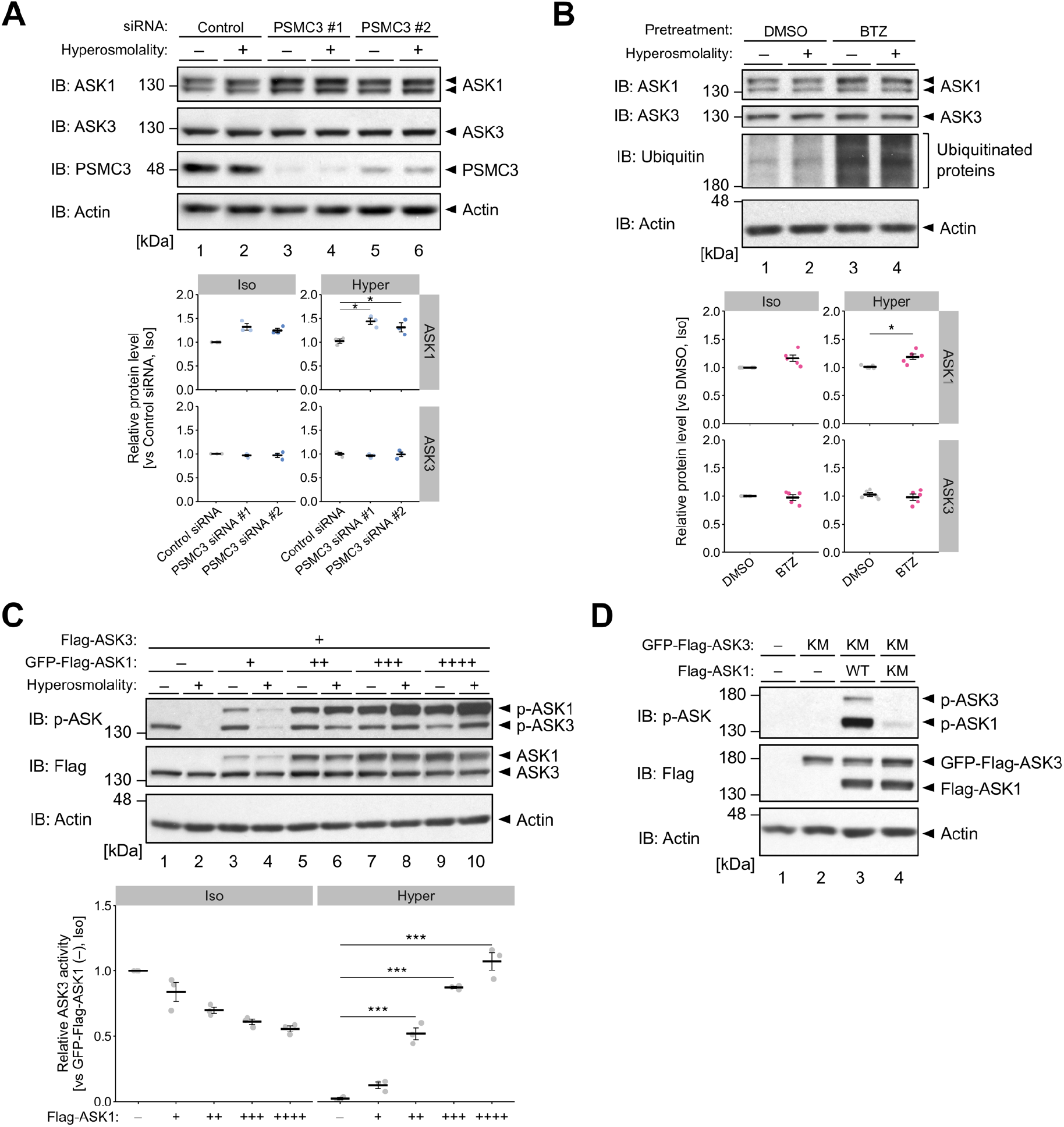
Increased ASK1 Protein Level Causes a Defect in ASK3 Inactivation under Proteasome Inhibition. (A) Effects of PSMC3 depletion on the protein levels of ASK1 and ASK3. (B) Effects of proteasome inhibition on the protein levels of ASK1 and ASK3. Cells were pretreated with 100 nM bortezomib for 6 h before osmotic stimuli. Proteasome inhibition is confirmed by the accumulation of ubiquitinated proteins. (C) Effects of ASK1 protein levels on ASK3 activity under hyperosmotic stress. Flag-ASK3 and GFP-Flag-ASK1 were exogenously expressed in cells. (D) Transphosphorylation of ASK3 by ASK1. GFP-Flag-ASK3 and Flag-ASK1 were exogenously expressed in cells. WT, wild-type; KM, kinase-negative mutant. (A–C) The bottom panels depict the quantification of western blots. Individual values and the mean ± SEM are presented as points and bars, respectively. *n* = 3 (A, C, D) and 5 (B). Iso, 300 mOsm (A–C); Hyper, 400 mOsm (A, B) or 500 mOsm (C); 10 min. IB, immunoblotting; BTZ, bortezomib. \**p* < 0.05, \*\*\**p* < 0.001. See also Figure S3.

It has been reported that ASK1 forms a heteromeric complex with another ASK family member, which decides basal ASK1 kinase activity via autophosphorylation and transphosphorylation ^23^. Given that ASK1 also forms a complex with ASK3 ^24^, we hypothesized that the proteasome inhibition increased the ASK1 protein level and the elevated ASK1 upregulated ASK3 activity via transphosphorylation under hyperosmotic stress. We therefore investigated the effect of ASK1 protein level on the activity of ASK3 under hyperosmotic stress. To control and evaluate the activities of ASK1 and ASK3 individually, we utilized green fluorescent protein (GFP) as a conjugating tag in an ASK1–ASK3 co-expression immunoblotting system, in which ASK3 was clearly distinguished from ASK1 by molecular weight. Increase in ASK1 protein level suppressed the ASK3 inactivation, and the highest condition of ASK1 co-expression even caused ASK3 activation under hyperosmotic stress (Figure 3C). This effect of ASK1 on ASK3 activity was dependent on the kinase activity of ASK1 because the phosphorylation on Thr808 of ASK3 kinase-negative mutant (KM), which loses the ability of autophosphorylation ^16^, was increased by co-expression of wild-type ASK1, but not by that of ASK1 KM ^25^ (Figure 3D). Therefore, these results suggest that the proteasome inhibits ASK1-mediated phosphorylation of ASK3 via degradation of ASK1 to ensure ASK3 inactivation under hyperosmotic stress.

### Proteasome Inhibition Disturbs ASK3 Signaling under Hyperosmotic Stress in an ASK1-Dependent Manner

Under hyperosmotic stress, ASK3 inactivation enables the activation of downstream kinase cascade including with-no-lysine (K) kinase 1 (WNK1), STE20/SPS1-related proline/alanine-rich kinase (SPAK), and oxidative stress-responsive kinase 1 (OSR1)^16^. We then examined whether the proteasomal regulation of ASK1 is involved in the ASK3-WNK1-SPAK/OSR1 signaling under hyperosmotic stress. ASK1 depletion tended to diminish the bortezomib-induced increase of ASK3 activity under hyperosmotic stress (Figure 4A; lanes 6 and 8), although ASK1 depletion reduced the basal ASK3 activity (lanes 5 and 7 in Fig. 4A) and thus potentially made the detection range too small to evaluate the effects of bortezomib on ASK3 activity. Additionally, the pretreatment of bortezomib inhibited hyperosmotic stress-induced SPAK/OSR1 activation in control cells but not in ASK1 knockdown cells (Figure 4A; lanes 2, 4, 6, and 8), suggesting that ASK1 mediates defect in the hyperosmotic stress-induced ASK3 signaling by proteasome inhibition. Moreover, ASK3 overexpression had much stronger inhibitory effect on SPAK/OSR1 activation under hyperosmotic stress than ASK1 overexpression (Figure 4B), further supporting that not the elevated ASK1 per se but the ASK1-transactivated ASK3 inhibited hyperosmotic stress-induced SPAK/OSR1 activation during proteasome inhibition. Given the potentially enormous effects of proteasome inhibition on broad spectrum of intracellular signaling, we alternatively induced accumulation of ASK1 by depletion of beta-transducin repeat containing protein 1 and 2 (β-TrCP1 and 2), E3 ubiquitin ligases for ASK1 ^26^. β-TrCP1/2 depletion increased ASK1 protein levels, but not ASK3 protein levels, and inhibited ASK inactivation and SPAK activation under hyperosmotic stress (Figure 4C; lanes 2 and 4). The effects of β-TrCP1/2 depletion on SPAK activation were abrogated by additional depletion of ASK1 (Figure 4C; lanes 4 and 8), suggesting that the elevated ASK1 protein level by β-TrCP1/2 depletion impaired SPAK activation under hyperosmotic stress. Collectively, these results suggest that the proteasome maintains ASK1 protein level to allow the ASK3 signaling under hyperosmotic stress.

**Figure 4.**
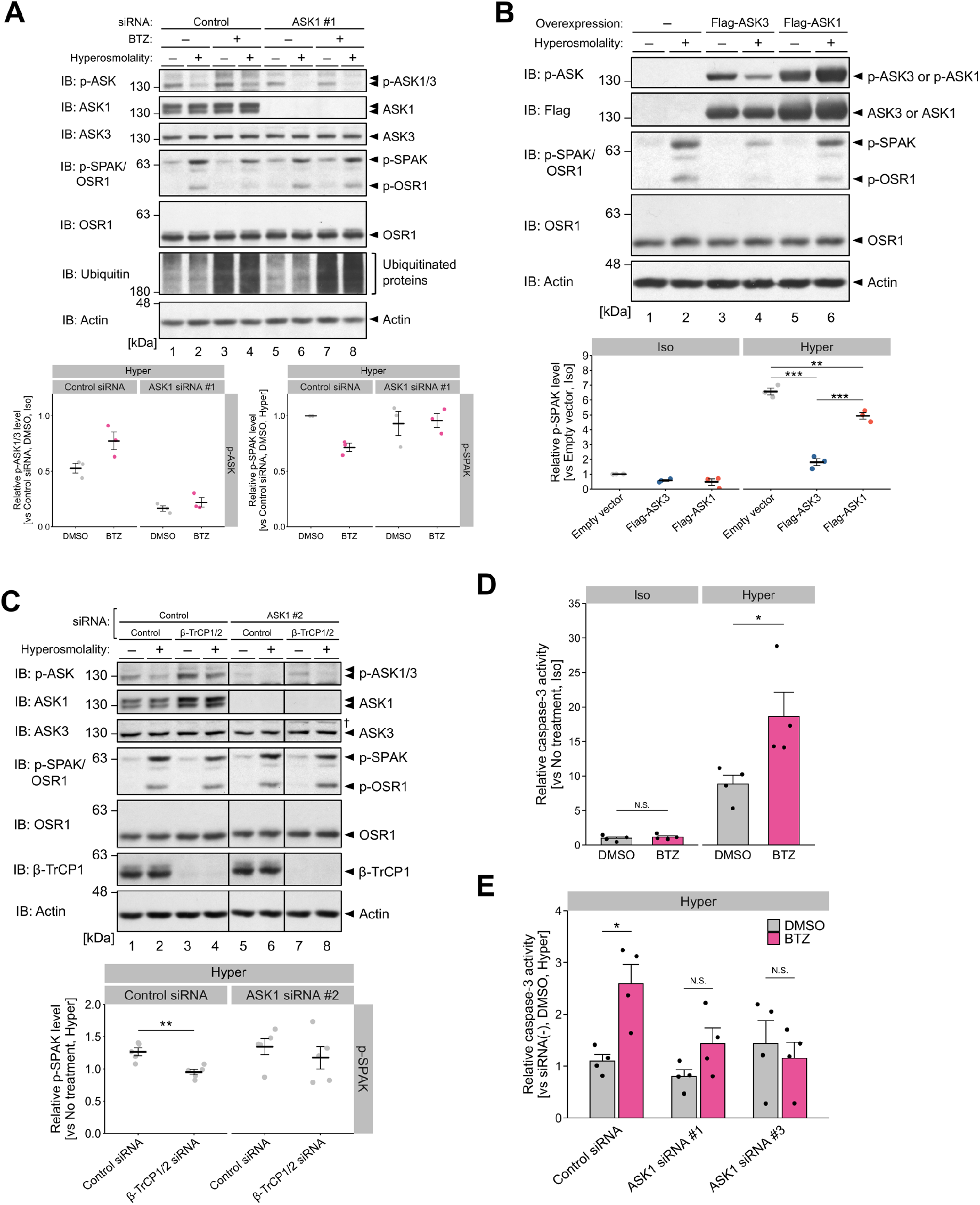
Proteasome Inhibition Causes Defects in ASK3 Signaling and Exacerbation of Apoptosis under Hyperosmotic Stress in an ASK1-Dependent Manner. (A) Requirement of ASK1 for the effects of proteasome inhibition on SPAK/OSR1 activation under hyperosmotic stress. Cells were pretreated with 100 nM bortezomib for 6 h before osmotic stimuli. (B) Effects of ASK1/3 overexpression on SPAK/OSR1 activity under hyperosmotic stress. (C) Requirement of ASK1 for the effects of β-TrCP1/2 depletion on SPAK/OSR1 activation under hyperosmotic stress. (D) Effects of proteasome inhibition on caspase-3 activation under hyperosmotic stress. (E) Effects of ASK1 depletion on bortezomib-induced overactivation of caspase-3 under hyperosmotic stress. (D, E) Cells were pretreated with 100 nM bortezomib for 6 h and the culture medium was changed to bortezomib-free osmotic medium. Mean + SEM. *n* = 4 (D, E). (A–C) The bottom panels depict the quantification of western blots. Individual values and the mean ± SEM are presented as points and bars, respectively. *n* = 3 (A, B) and 5 (C). Iso, 300 mOsm (A–C) or culture medium (D); Hyper, 400 mOsm (A– C) culture medium supplemented with 500 mM mannitol (D, E); 10 min (A–C) or 12 h (D, E). BTZ, bortezomib. N.S., not significant; \**p* < 0.05, \*\**p* < 0.01, \*\*\**p* < 0.0001.

### Proteasome inhibition causes ASK1-mediated exacerbation of apoptosis under hyperosmotic stress

Since ASK1 is well known to induce apoptosis under various stresses ^27^, we hypothesized that the proteasomal maintenance of ASK1 protein level is a mechanism linking the proteostasis overload and cell fate under hyperosmotic stress. We first tested if disruption of proteasome function decreases the survival of mammalian cells under hyperosmotic stress. We pretreated cells with bortezomib and changed the culture medium to either isoosmotic or hyperosmotic buffer without bortezomib. This transient bortezomib pretreatment did not activate caspase-3 under isoosmotic buffer but accelerated caspase-3 activation under hyperosmotic stress (Figure 4D), showing that the precondition of proteasome inhibition made cells prone to undergo hyperosmotic stress-induced apoptosis. We then tested if ASK1 is required for the exacerbation of hyperosmotic stress-induced apoptosis by the proteotoxic preconditioning. ASK1 knockdown reduced the effect of proteasome inhibition on caspase-3 activity under hyperosmotic stress (Figure 4E). Therefore, proteasome inhibition accelerates apoptosis under hyperosmotic stress in an ASK1-dependent manner.

## Discussion

Although the importance of proteostasis maintenance in the survival of *C*.*elegans* and mammalian cells under hyperosmotic stress has been reported ^13–15^, it has been unknown whether particular molecules sense the integrity of cellular proteostasis pathways to regulate cell survival and death responses under hyperosmotic stress. Here, we demonstrated that insufficient proteasome capacity led to the accumulation of ASK1 under hyperosmotic stress, leading to the failure of ASK3 signaling and the aggravation of hyperosmotic stress-induced apoptosis (Figures 3 and 4). Given that the proteasome is one of the major degradation machineries for damaged proteins, ASK1-dependent regulation of apoptosis can contribute to the elimination of the cells which are prone to retain damaged proteins due to low proteasome activity under hyperosmotic stress. Proteasome inhibition also sensitizes cells to other proteotoxic stresses such as oxidative stress and heat shock via unknown mechanisms ^28–30^. Degradation of ASK1 by the ubiquitin-proteasome system has been reported to suppress cell death under oxidative stress ^26,31–33^. Therefore, it is possible that the proteasomal maintenance of ASK1 protein level underlies the bifurcation between cell survival and death responses in diverse proteotoxic conditions beyond hyperosmotic stress.

In addition to the proteasomal maintenance of ASK1, the intra-relationship of ASK family proteins should be highlighted for the apoptosis-inducing function of ASK1. Elevated ASK1 protein level enhanced ASK1-mediated transphosphorylation to ASK3, thereby inhibiting ASK3 inactivation under hyperosmotic stress (Figures 3C and 3D). Interestingly, our results suggested that activity regulation between ASK1 and ASK3 is reciprocal. In contrast to ASK3, ASK1 is activated in response to hyperosmotic stress^16^ (Figure 4B; lanes 5 and 6).This ASK1 activation was attenuated as the relative protein level of ASK1 to ASK3 decreased (Figure 3C; lanes 5–10), and even ASK1 inactivation was observed in the lowest ASK1-to-ASK3 ratio condition (Figure 3C; lanes 3 and 4). In addition, the overall activity of endogenous ASK1 and ASK3 was decreased upon hyperosmotic stress (e.g., lanes 1 and 2 in Figure 1F). These results indicate that ASK3 negatively regulates ASK1 activity under hyperosmotic stress. Knockdown of ASK3 tended to further enhance the ASK1-dependent apoptosis of bortezomib-pretreated cells under hyperosmotic stress (Figure S4A), which may be due to overactivation of ASK1 as a result of its release from negative regulation by ASK3. Collectively, we propose that the protein level ratio of ASK1 to ASK3 has a key role as an index of the integrity of proteasome activity to decide the bifurcation between cellular survival and death responses under hyperosmotic stress. ASK3 is predominantly expressed in tissues that face extracellular fluid possessing broad range of osmolality such as kidneys and stomach, while ASK1 is ubiquitously expressed ^16,34^. Therefore, cells in the kidney and stomach may acquire higher resistance to hyperosmotic stress-induced apoptosis by low ASK1-to-ASK3 expression ratio. In mouse models of kidney disease, it has been reported that ASK1 is activated and mediates the progression or aggravation of pathological conditions ^35,36^. It is of great interest to examine whether the ASK1-to-ASK3 ratio controls any physiological function of ASK1 in kidney disease.

The physiological consequences of the proteasomal regulation of hyperosmotic stress-induced ASK3 inactivation remain unclear. Given that ASK3 inactivation has an essential role in RVI under hyperosmotic stress ^17^ and that inhibition of RVI is a prerequisite for hyperosmotic stress-induced apoptosis ^37^, we assumed that insufficient ASK3 inactivation by proteasome inhibition would promote apoptosis under hyperosmotic stress. However, as mentioned above, depletion of ASK3 did not attenuate but rather tended to accelerate apoptosis under hyperosmotic stress in proteasome-inhibited cells (Figure S4A), suggesting a central role of ASK1 but not ASK3 as the “judge” of hyperosmotic stress-induced apoptosis execution for cells with low proteostasis capacity. ASK3 inhibited hyperosmotic stress-induced SPAK/OSR1 activation when ASK1 protein level was increased by β-TrCP1/2 depletion (Figure S4B), but the downstream signaling effects on cellular events should be investigated in future studies.

In summary, we revealed that proteasomal regulation of ASK1 protein level is important for the modulation of ASK3 signaling and apoptosis under hyperosmotic stress. Our findings emphasize the role of proteasome activity and expression ratio of ASK1 and ASK3 in the determination of cell fate bifurcation under, and possibly beyond, hyperosmotic stress.

## Materials and Methods

### RESOURCE AVAILABILITY

#### Lead Contact

Further information and requests for the resources and reagents should be directed to the lead contact, Hidenori Ichijo.

#### Materials availability

Reagents generated in this study will be made available on request.

#### Data and code availability

All data reported in this study will be shared by the lead contact upon request. This paper does not report original code.

Any additional information required to reanalyze the data reported in this work is available from the lead contact upon request.

### EXPERIMENTAL MODEL AND SUBJECT DETAILS

#### Cell lines and cell culture

HEK293A cells were cultured in Dulbecco’s Modified Eagle’s Medium (DMEM)-high glucose (Sigma-Aldrich, Cat#D5796) supplemented with 10% FBS and 100 units/mL penicillin G (Meiji Seika, Cat#6111400D2039). Tetracycline-inducible Flag-ASK3-stably-expressing HEK293A cells^17^ were cultured in DMEM-high glucose supplemented with 10% FBS, 2.5 μg/mL blasticidin (Invitrogen, Cat#46-1120 or Cat#A1113903) and 50 μg/mL Zeocin (Invitrogen, Cat#R25001). To induce Flag-ASK3, cells were pre-treated with 1 µg/mL tetracycline (Sigma-Aldrich, Cat#T7660) before assays. All cells were cultured in 5% CO_2_ at 37°C and verified to be negative for mycoplasma.

### METHOD DETAILS

#### Functional enrichment analysis

Metascape^38^ (https://metascape.org) was used to perform the Gene Ontology enrichment analysis and the protein-protein interaction (PPI) analysis on the positive genes identified in the genome-wide siRNA screen for the regulators of ASK3 inactivation under hyperosmotic stress^17^. 63 positive genes in the second screen were submitted for input and the analysis was run with default settings except for background genes, which were set to the genes targeted by the genome-wide siRNA library. The Gene Ontology enrichment analysis was carried out using the following ontology sources: KEGG Pathway, GO Biological Processes, Reactome Gene Sets, Canonical Pathways, Cell Type Signatures, CORUM, TRRUST, DisGeNET, PaGenBase, Transcription Factor Targets, WikiPathways and COVID. The PPI analysis was carried out using the following databases: STRING, BioGrid, OmniPath, and InWeb_IM. Only physical interactions in STRING (physical score > 0.132) and BioGrid were used.

#### Transfections

Plasmid transfections were performed with polyethylenimine “MAX” (Polysciences, Cat#24765) when HEK293A cells were grown to 60–90% confluency, according to a previously described protocol^39^ with minor optimization. siRNA transfections for cells at 10–40% confluency were carried out by forward (for Figure 4C, S1D, and S4B) or reverse (for other figures) transfection using Lipofectamine™ RNAiMAX (Invitrogen, Cat#13778-150) at a final concentration of 5–30 nM for 48 h, according to the manufacturer’s instructions. For experiments in which both plasmid and siRNA were transfected, siRNA transfections were performed 24 h before plasmid transfections.

#### Expression plasmids

Expression plasmids for this study were constructed by standard molecular biology techniques and all constructs were verified by sequencing. As for the EGFP-Flag-ASK1 expression plasmid, ASK1 cDNA^23^ was subcloned into pcDNA3/GW with an N-terminal EGFP-Flag-tag^18^.

#### Osmotic stress treatments

Osmotic stress was applied by exchanging culture medium for osmotic buffers. Isoosmotic buffer (300 mOsm/kg H_2_O, pH 7.4) contained 130 mM NaCl, 2 mM KCl, 1 mM KH_2_PO_4_, 2 mM CaCl_2_, 2 mM MgCl_2_, 10 mM HEPES, 10 mM glucose, and 20 mM mannitol. Hyperosmotic buffer (400 or 500 mOsm/kg H_2_O, pH 7.4) was the same as isoosmotic buffer but contained 120 or 220 mM mannitol, respectively. The 400 or 500 mOsm/kg H_2_O hyperosmotic buffers were created by mixing isoosmotic buffer and 800 mOsm/kg H_2_O hyperosmotic buffer. Absolute osmolality was verified by Osmomat 030 (Gonotec) to fall within a range from 295 to 315 mOsm for isoosmotic buffer and ± 25 mOsm/kg H_2_O for the other buffers. In Figures 4D–E, S3A, and S4A, osmotic stress treatment was applied by changing to fresh culture medium supplemented with mannitol and incubating in 5% CO_2_ at 37°C. For proteasome inhibition, bortezomib (LC Laboratories, Cat#B-1408) or MG132 (Enzo Life Sciences, Cat#BML-PI102) dissolved in DMSO was added to culture medium before osmotic stress treatments. For translation inhibition, cycloheximide (Sigma-Aldrich, Cat#C7698-1G) dissolved in DMSO was added to culture medium 2 h before osmotic stress treatments.

#### Cell lysis and western blotting

Cells were lysed in a lysis buffer (20 mM Tris-HCl pH 7.5, 150 mM NaCl, 10 mM EDTA, 1% sodium deoxycholate, and 1% Triton X-100) supplemented with 1 mM phenylmethylsulfonyl fluoride (PMSF), 5 µg/mL leupeptin, and phosphatase inhibitor cocktail (20 mM NaF, 30 mM b-glycerophosphate, 2.5 mM Na_3_VO_4_, 3 mM Na_2_MoO_4_, 12.5 µM cantharidin, and 5 mM imidazole). Cell extracts were clarified by centrifugation, and the supernatant was sampled by adding an equal volume of 2x SDS sample buffer (80 mM Tris HCl pH 8.8, 80 µg/mL bromophenol blue, 28.8% glycerol, 4% SDS, and 10 mM dithiothreitol). After boiling at 98°C for 3 min, the samples were resolved by SDS-PAGE and electroblotted onto a BioTrace PVDF membrane (Pall), FluoroTrans W membrane (Pall), or Immobilon-P membrane (Millipore, Cat#IPVH00010). The membranes were blocked with 3% or 5% skim milk (Megmilk Snow Brand) in TBS-T (20 mM Tris-HCl pH8.0, 137 mM NaCl and 0.1% Tween 20) and probed with the appropriate primary antibodies diluted by 1^st^ antibody-dilution buffer (TBS-T supplemented with 5% BSA (Iwai Chemicals, Cat#A001) and 0.1% NaN_3_ (Nacalai Tesque, Cat#312-33)). After replacing and probing the appropriate secondary antibodies diluted by TBS-T containing 3% or 5% skim milk, antibody-antigen complexes were detected on X-ray films (FUJIFILM, Cat#47410-07523, Cat#47410-26615 or Cat#47410-07595) or with a FUSION Solo S imaging system (VILBER) using ECL reagents (GE Healthcare). Quantification was performed via densitometry using ImageJ^40^. ASK3 activity was defined as the ratio of phosphorylated protein to total protein. Representative images were adjusted to the appropriate brightness and contrast using the GNU Image Manipulation Program (GIMP).

#### Immunofluorescence-based assay system for ASK3 activity

Tetracycline-inducible FLAG-ASK3-stably expressing HEK293A cells were seeded in 96-well plate. The cells were transfected with siRNA or treated with proteasome inhibitors followed by osmotic stress for 10 min. Quantification of ASK3 activity was performed using the immunofluorescence-based high content analysis reported from our group ^16,17^ with minor optimization. Immunostaining was performed by following procedure: fixation for 15 min with 4% formaldehyde in PBS, permeabilization for 15 min with 0.2% Triton X-100 in PBS, blocking for 30 min with 5% skim milk in TBS-T, and incubation at 4°C overnight with the primary antibodies in antibody-dilution buffer. The cells were further incubated at room temperature in the dark for 1 h with the appropriate fluorophore-conjugated secondary antibodies in TBS-T. After staining with Hoechst 33258 dye (Dojindo, Cat#343-07961, 1:1,000) in TBS-T for 5 min, the ASK3 activity was measured and analyzed by using CellInsight NXT (Thermo Fisher Scientific) with HCS Studio (Thermo Fisher Scientific). The white point, black rectangle, and black line in each violin plot indicate the median, interquartile range (IQR), and the range from upper quartile plus 1.5xIQR to lower quartile minus 1.5xIQR, respectively. The area of the violin plot is drawn to be equal across all experimental conditions.

#### Quantitative RT-PCR

Cells were seeded in 12-well plate. Two days after seeding, cells were treated with proteasome inhibitors when cell density reached 50–80%. Total RNA was isolated from cells using Isogen (Wako, Cat#319-90211) and reverse transcribed with ReverTra Ace qPCR RT Master Mix with gDNA Remover (Toyobo, Cat#FSQ-301), according to manufacturer’s instruction. Quantitative PCR was carried out using a LightCycler 96 (Roche) using Kapa SYBR Fast qPCR Master Mix (Kapa Biosystems, Cat#KK4602). Data were normalized to GAPDH, and primer sequences are listed in Table S1.

#### Caspase-3 activity assay

Apoptosis induction was measured using a fluorogenic substrate for activated caspase-3, Ac-DEVD-AFC (Cayman, Cat#14459). Cells were lysed with PBS containing 0.1 % Triton X-100 after osmotic stress, and the cell lysate was centrifuged at 17,700 × g, 4°C for 10 min. Lysate samples were individually mixed with reagents in 96-well microplates (25 μL of lysate sample, 49.5 μL of 2 × Reaction Buffer (BioVision, Cat#1068), 25 μL of PBS, 5 μL of caspase-3 substrate (1 mM in DMSO) and 0.5 μL of 1 M dithiothreitol (TCI, Cat#D1071)). Fluorescence signals were measured at specific wavelengths (Ex/Em = 400/505 nm) using a Varioskan Flash (Thermo Fisher Scientific) after incubation at 37 °C for approximately 90 min. For normalization, the protein amount in each lysate sample was measured using a DC™ protein assay (Bio-Rad, Cat#5000113, #5000114, #5000115). Caspase-3 activity in each sample was calculated as follows: ((fluorescence intensity of sample − background)/protein concentration of sample (μg/μL)). Relative Caspase-3 activity was determined by standardizing the caspase-3 activity of each sample by the caspase-3 activity under the conditions shown in figure.

#### Proteasome activity assay

Cells were seeded in 24-well plate. Two days after seeding, cells were treated with proteasome inhibitors when cell density reached 70–90%. Cells were lysed in ice-cold buffer containing 25 mM Tris-HCl (pH 7.5), 0.2% Nonidet P-40, 1 mM dithiothreitol, 2 mM ATP, and 5 mM MgCl_2_. The lysate was mixed with the fluorogenic peptide, succinyl-Leu-Leu-Val-Tyr-7-amino-4-methylcoumarin (Suc-LLVY-MCA; Peptide Institute, Cat#3120-v) in 100 mM Tris-HCl (pH 8.0) at 37°C. The hydrolysis of Suc-LLVY-MCA was measured by Varioskan Flash (Ex/Em = 360/460 nm).

### QUANTIFICATION AND STATISTICAL ANALYSIS

The statistical results are expressed as the mean ± SEM unless otherwise indicated. No statistical method was utilized to predetermine sample size. In the case of ASK3 activity determined by quantification of immunofluorescence, Wilcoxon rank sum test was used for comparisons between two groups, and Kruskal-Wallis test followed by Dunn’s test was used for comparisons between three groups. For other parameters, unpaired two-tailed Student’s *t*-test was used for comparisons between two groups, and Dunnett’s test or Tukey-Kramer’s test was used for comparisons between three or more groups. Number of samples and sample size are indicated in the figure legends and no sample was excluded from statistical tests. Statistical tests were performed using GraphPad Prism version 7.0c for Mac OS X, GraphPad Software, San Diego, California USA, www.graphpad.com. Other statistical tests were performed using R with RStudio (RStudio, Inc., Boston, MA. https://www.rstudio.com). *P* < 0.05 was considered statistically significant. The investigators were not blinded to allocation during experiments and outcome assessment. The experiments were not randomized.

## Acknowledgments

We thank all members of the Laboratory of Cell Signaling for meaningful discussion. This work was supported by the Japan Agency for Medical Research and Development (AMED) under the Project for Elucidating and Controlling Mechanisms of Aging and Longevity (grant number JP21gm5010001 to H.I.), Japan Society for the Promotion of Science (JSPS) under the Grants-in-Aid for Scientific Research (KAKENHI; JP17J10998 to X.Z., JP19K16067 to K.W., JP2304918 to S.M., JP18H02569 and JP22K19324 to I.N., and JP18H03995 and JP21H04760 to H.I.), Japan Science and Technology Agency (JST) Moonshot R&D—MILLENNIA Program (grant number JPMJMS2022-18 to H.I.), and by AMED Project for LEAP (grant number 23gm0010009s0102 to H.I.).

## Author Contributions

X.Z., K.W., I.N., and H.I. conceptualized this study. X.Z. designed and performed almost all experiments. K.M. performed the ASK1–ASK3 co-expression experiments. J.H. and S.M. provided reagents and expertise. X.Z. statistically analyzed and visualized the data. K.W., I.N., and H.I. supervised this project. X.Z., K.W., I.N., and H.I. wrote the manuscript.

## Declaration of Interests

The authors declare no competing interests.

## Supplemental Information

Supplemental Information includes four figures and two tables. Document S1. Supplemental Figures S1–S4 and Table S1 and S2

## Notes

### Competing Interest Statement

The authors have declared no competing interest.

## References

1. King, L.S., Kozono, D., and Agre, P. (2004). From structure to disease: the evolving tale of aquaporin biology. Nat. Rev. Mol. Cell Biol. 5, 687–698. 10.1038/nrm1469.

2. Brocker, C., Thompson, D.C., and Vasiliou, V. (2012). The role of hyperosmotic stress in inflammation and disease. Biomol. Concepts 3, 345–364. 10.1515/bmc-2012-0001.

3. Burg, M.B., Ferraris, J.D., and Dmitrieva, N.I. (2007). Cellular response to hyperosmotic stresses. Physiol. Rev. 87, 1441–1474. 10.1152/physrev.00056.2006.

4. Hoffmann, E.K., Lambert, I.H., and Pedersen, S.F. (2009). Physiology of cell volume regulation in vertebrates. Physiol. Rev. 89, 193–277. 10.1152/physrev.00037.2007.

5. Dominy, B.N., Perl, D., Schmid, F.X., and Brooks, C.L., 3rd (2002). The effects of ionic strength on protein stability: the cold shock protein family. J. Mol. Biol. 319, 541–554. 10.1016/S0022-2836(02)00259-0.

6. Kohn, W.D., Kay, C.M., and Hodges, R.S. (1997). Salt effects on protein stability: two-strandedα-helical coiled-coils containing inter-or intrahelical ion pairs. J. Mol. Biol. 267, 1039–1052.

7. Yancey, P.H., Clark, M.E., Hand, S.C., Bowlus, R.D., and Somero, G.N. (1982). Living with water stress: evolution of osmolyte systems. Science 217, 1214–1222. 10.1126/science.7112124.

8. Ellis, R.J., and Minton, A.P. (2006). Protein aggregation in crowded environments. Biol. Chem. 387, 485–497. 10.1515/BC.2006.064.

9. Munishkina, L.A., Ahmad, A., Fink, A.L., and Uversky, V.N. (2008). Guiding protein aggregation with macromolecular crowding. Biochemistry 47, 8993–9006. 10.1021/bi8008399.

10. Zhou, H.-X., Rivas, G., and Minton, A.P. (2008). Macromolecular crowding and confinement: biochemical, biophysical, and potential physiological consequences. Annu. Rev. Biophys. 37, 375–397. 10.1146/annurev.biophys.37.032807.125817.

11. Burkewitz, K., Choe, K., and Strange, K. (2011). Hypertonic stress induces rapid and widespread protein damage in C. elegans. Am. J. Physiol. Cell Physiol. 301, C566–76. 10.1152/ajpcell.00030.2011.

12. Mazzeo, L.E.M., Dersh, D., Boccitto, M., Kalb, R.G., and Lamitina, T. (2012). Stress and aging induce distinct polyQ protein aggregation states. Proceedings of the National Academy of Sciences 109, 10587–10592.

13. Choe, K.P., and Strange, K. (2008). Genome-wide RNAi screen and in vivo protein aggregation reporters identify degradation of damaged proteins as an essential hypertonic stress response. Am. J. Physiol. Cell Physiol. 295, C1488–98. 10.1152/ajpcell.00450.2008.

14. Yasuda, S., Tsuchiya, H., Kaiho, A., Guo, Q., Ikeuchi, K., Endo, A., Arai, N., Ohtake, F., Murata, S., Inada, T., et al. (2020). Stress- and ubiquitylation-dependent phase separation of the proteasome. Nature 578, 296–300. 10.1038/s41586-020-1982-9.

15. Nunes, P., Ernandez, T., Roth, I., Qiao, X., Strebel, D., Bouley, R., Charollais, A., Ramadori, P., Foti, M., Meda, P., et al. (2013). Hypertonic stress promotes autophagy and microtubule-dependent autophagosomal clusters. Autophagy 9, 550–567.

16. Naguro, I., Umeda, T., Kobayashi, Y., Maruyama, J., Hattori, K., Shimizu, Y., Kataoka, K., Kim-Mitsuyama, S., Uchida, S., Vandewalle, A., et al. (2012). ASK3 responds to osmotic stress and regulates blood pressure by suppressing WNK1-SPAK/OSR1 signaling in the kidney. Nat. Commun. 3, 1285. 10.1038/ncomms2283.

17. Watanabe, K., Umeda, T., Niwa, K., Naguro, I., and Ichijo, H. (2018). A PP6-ASK3 Module Coordinates the Bidirectional Cell Volume Regulation under Osmotic Stress. Cell Rep. 22, 2809–2817. 10.1016/j.celrep.2018.02.045.

18. Watanabe, K., Morishita, K., Zhou, X., Shiizaki, S., Uchiyama, Y., Koike, M., Naguro, I., and Ichijo, H. (2021). Cells recognize osmotic stress through liquid-liquid phase separation lubricated with poly(ADP-ribose). Nat. Commun. 12, 1353. 10.1038/s41467-021-21614-5.

19. Ichijo, H., Nishida, E., Irie, K., ten Dijke, P., Saitoh, M., Moriguchi, T., Takagi, M., Matsumoto, K., Miyazono, K., and Gotoh, Y. (1997). Induction of apoptosis by ASK1, a mammalian MAPKKK that activates SAPK/JNK and p38 signaling pathways. Science 275, 90–94. 10.1126/science.275.5296.90.

20. Tobiume, K., Saitoh, M., and Ichijo, H. (2002). Activation of apoptosis signal-regulating kinase 1 by the stress-induced activating phosphorylation of pre-formed oligomer. Journal of Cellular Physiology 191, 95–104. 10.1002/jcp.10080.

21. Murata, S., Yashiroda, H., and Tanaka, K. (2009). Molecular mechanisms of proteasome assembly. Nat. Rev. Mol. Cell Biol. 10, 104–115. 10.1038/nrm2630.

22. Sherman, D.J., and Li, J. (2020). Proteasome Inhibitors: Harnessing Proteostasis to Combat Disease. Molecules 25. 10.3390/molecules25030671.

23. Takeda, K., Shimozono, R., Noguchi, T., Umeda, T., Morimoto, Y., Naguro, I., Tobiume, K., Saitoh, M., Matsuzawa, A., and Ichijo, H. (2007). Apoptosis signal-regulating kinase (ASK) 2 functions as a mitogen-activated protein kinase kinase kinase in a heteromeric complex with ASK1. J. Biol. Chem. 282, 7522–7531. 10.1074/jbc.M607177200.

24. Federspiel, J.D., Codreanu, S.G., Palubinsky, A.M., Winland, A.J., Betanzos, C.M., McLaughlin, B., and Liebler, D.C. (2016). Assembly Dynamics and Stoichiometry of the Apoptosis Signal-regulating Kinase (ASK) Signalosome in Response to Electrophile Stress. Mol. Cell. Proteomics 15, 1947–1961. 10.1074/mcp.M115.057364.

25. Chang, H.Y., Nishitoh, H., Yang, X., Ichijo, H., and Baltimore, D. (1998). Activation of apoptosis signal-regulating kinase 1 (ASK1) by the adapter protein Daxx. Science 281, 1860–1863. 10.1126/science.281.5384.1860.

26. Cheng, R., Takeda, K., Naguro, I., Hatta, T., Iemura, S.-I., Natsume, T., Ichijo, H., and Hattori, K. (2018). β-TrCP-dependent degradation of ASK1 suppresses the induction of the apoptotic response by oxidative stress. Biochim. Biophys. Acta Gen. Subj. 1862, 2271–2280. 10.1016/j.bbagen.2018.07.015.

27. Nishida, T., Hattori, K., and Watanabe, K. (2017). The regulatory and signaling mechanisms of the ASK family. Adv. Biol. Regul. 66, 2–22. 10.1016/j.jbior.2017.05.004.

28. Ding, Q., and Keller, J.N. (2001). Proteasome inhibition in oxidative stress neurotoxicity: implications for heat shock proteins. J. Neurochem. 77, 1010–1017. 10.1046/j.1471-4159.2001.00302.x.

29. Mytilineou, C., McNaught, K.S.P., Shashidharan, P., Yabut, J., Baptiste, R.J., Parnandi, A., and Olanow, C.W. (2004). Inhibition of proteasome activity sensitizes dopamine neurons to protein alterations and oxidative stress. J. Neural Transm. 111, 1237–1251. 10.1007/s00702-004-0167-2.

30. Alvarez-Berríos, M.P., Castillo, A., Rinaldi, C., and Torres-Lugo, M. (2014). Magnetic fluid hyperthermia enhances cytotoxicity of bortezomib in sensitive and resistant cancer cell lines. Int. J. Nanomedicine 9, 145–153. 10.2147/IJN.S51435.

31. Maruyama, T., Araki, T., Kawarazaki, Y., Naguro, I., Heynen, S., Aza-Blanc, P., Ronai, Z., Matsuzawa, A., and Ichijo, H. (2014). Roquin-2 promotes ubiquitin-mediated degradation of ASK1 to regulate stress responses. Sci. Signal. 7, ra8. 10.1126/scisignal.2004822.

32. Zhang, Z., Hao, J., Zhao, Z., Ben, P., Fang, F., Shi, L., Gao, Y., Liu, J., Wen, C., Luo, L., et al. (2009). beta-Arrestins facilitate ubiquitin-dependent degradation of apoptosis signal-regulating kinase 1 (ASK1) and attenuate H2O2-induced apoptosis. Cell. Signal. 21, 1195–1206. 10.1016/j.cellsig.2009.03.010.

33. Nagai, H., Noguchi, T., Homma, K., Katagiri, K., Takeda, K., Matsuzawa, A., and Ichijo, H. (2009). Ubiquitin-like sequence in ASK1 plays critical roles in the recognition and stabilization by USP9X and oxidative stress-induced cell death. Mol. Cell 36, 805–818. 10.1016/j.molcel.2009.10.016.

34. Iriyama, T., Takeda, K., Nakamura, H., Morimoto, Y., Kuroiwa, T., Mizukami, J., Umeda, T., Noguchi, T., Naguro, I., Nishitoh, H., et al. (2009). ASK1 and ASK2 differentially regulate the counteracting roles of apoptosis and inflammation in tumorigenesis. EMBO J. 28, 843–853. 10.1038/emboj.2009.32.

35. Liles, J.T., Corkey, B.K., Notte, G.T., Budas, G.R., Lansdon, E.B., Hinojosa-Kirschenbaum, F., Badal, S.S., Lee, M., Schultz, B.E., Wise, S., et al. (2018). ASK1 contributes to fibrosis and dysfunction in models of kidney disease. J. Clin. Invest. 128, 4485–4500. 10.1172/JCI99768.

36. Ma, F.Y., Tesch, G.H., and Nikolic-Paterson, D.J. (2014). ASK1/p38 signaling in renal tubular epithelial cells promotes renal fibrosis in the mouse obstructed kidney. Am. J. Physiol. Renal Physiol. 307, F1263–73. 10.1152/ajprenal.00211.2014.

37. Maeno, E., Ishizaki, Y., Kanaseki, T., Hazama, A., and Okada, Y. (2000). Normotonic cell shrinkage because of disordered volume regulation is an early prerequisite to apoptosis. Proc. Natl. Acad. Sci. U. S. A. 97, 9487–9492. 10.1073/pnas.140216197.

38. Zhou, Y., Zhou, B., Pache, L., Chang, M., Khodabakhshi, A.H., Tanaseichuk, O., Benner, C., and Chanda, S.K. (2019). Metascape provides a biologist-oriented resource for the analysis of systems-level datasets. Nat. Commun. 10, 1523. 10.1038/s41467-019-09234-6.

39. Longo, P.A., Kavran, J.M., Kim, M.-S., and Leahy, D.J. (2013). Transient mammalian cell transfection with polyethylenimine (PEI). Methods Enzymol. 529, 227–240. 10.1016/B978-0-12-418687-3.00018-5.

40. Schneider, C.A., Rasband, W.S., and Eliceiri, K.W. (2012). NIH Image to ImageJ: 25 years of image analysis. Nat. Methods 9, 671–675. 10.1038/nmeth.2089.

