## Supplementary Material for "Proteasomal regulation of ASK family kinases dictates cell fate under hyperosmotic stress"

**This PDF file includes:**

Figure S1 to S4

Table S1 and S2

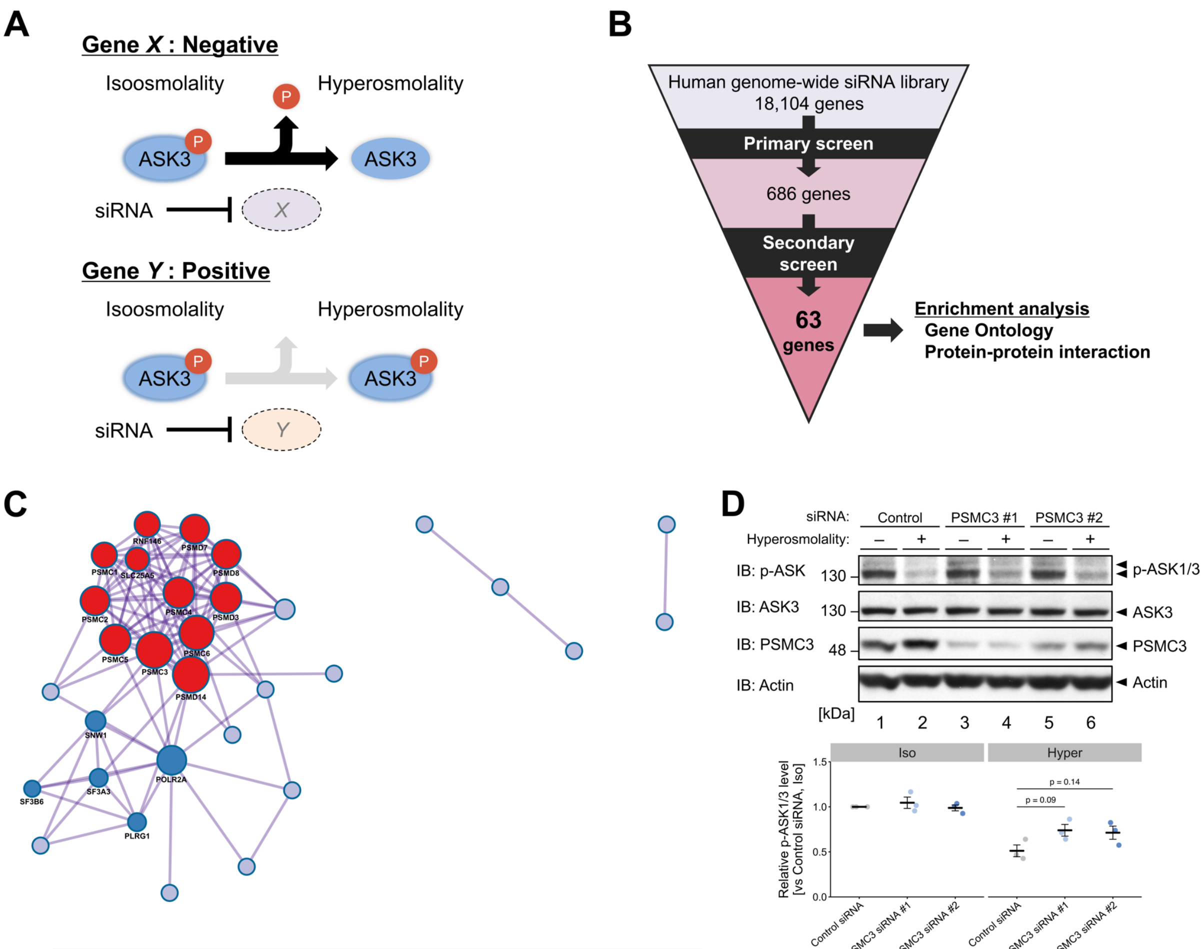


| Color | MCODE | Annotation ID | Description | Log10(P) |
|---|---|---|---|---|
| (red) | MCODE_1 | CORUM:32 | PA700 complex | –28.9 |
| (red) | MCODE_1 | R-HSA-4641257 | Degradation of AXIN | –27.1 |
| (red) | MCODE_1 | R-HSA-8948751 | Regulation of PTEN stability and activity | –26.0 |
| (blue) | MCODE_2 | R-HSA-72163 | mRNA Splicing – Major Pathway | –10.0 |
| (blue) | MCODE_2 | R-HSA-72172 | mRNA Splicing | –9.9 |
| (blue) | MCODE_2 | R-HSA-72203 | Processing of Capped Intron-Containing Pre-mRNA | –9.4 |

**Figure S1. The proteasome as a negative regulator of ASK3 activity under hyperosmotic stress. Related to Figure 1.**

(A, B) Rationale (A) and process (B) of the siRNA screen for the regulators of ASK3 inactivation under hyperosmotic stress, related to Figure 1A.

(C) Protein–protein interaction (PPI) networks and Molecular Complex Detection (MCODE; [1]) components identified in the positive genes of the secondary screen, related to Figure 1C. Nodes indicate proteins that physically interact with at least one of the other proteins encoded by the positive genes, and edges indicate their PPIs. The densely connected clusters of proteins identified by application of the MCODE algorithm to the PPI network are shown in red or blue. Pathway and process enrichment analysis was applied to each MCODE component, and the three best-scoring terms by nominal *p*-value are shown in the lower panel.

(D) Effects of PSMC3 depletion on endogenous ASK activity under hyperosmotic stress, related to Figure 1F. The bottom panel depicts the quantification of western blots. Individual values and the mean ± SEM are presented as points and bars, respectively. *n* = 3. Iso, 300 mOsm; Hyper, 400 mOsm; 10 min. IB, immunoblotting.

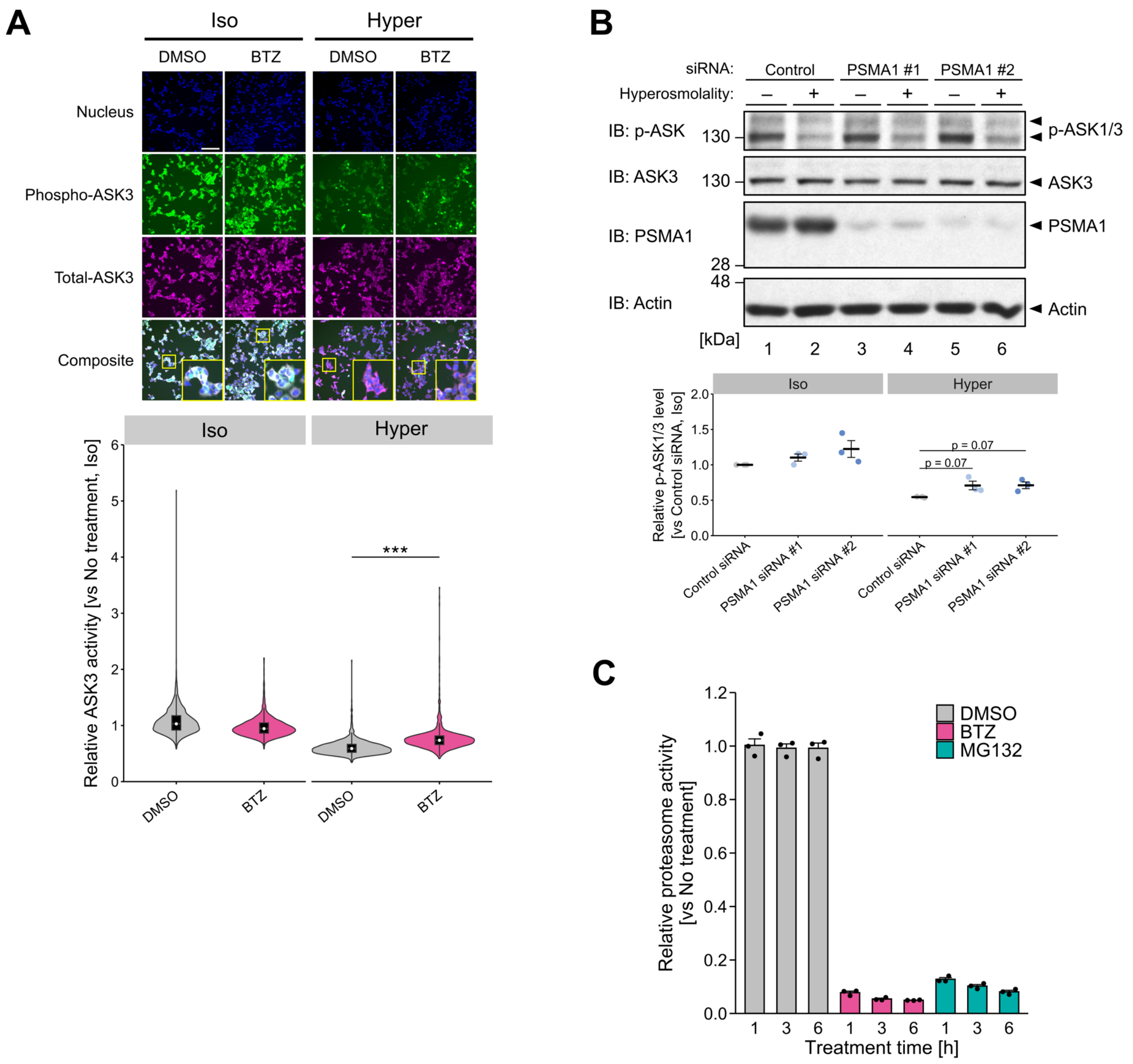


**Figure S2. Requirement of proteasome activity for ASK3 inactivation under hyperosmotic stress. Related to Figure 2.**

(A) Effects of bortezomib on ASK3 activity in tetracycline-inducible Flag-ASK3-stably expressing cells, related to Figure 2A. Cells were pretreated with 100 nM bortezomib for 6 h before osmotic stimuli. ASK3 activity was measured as in Figure 1D. The top panel shows immunofluorescence images of nuclei, phospho-ASK3, total-ASK3, and their composite. The bottom panel shows violin plots of ASK3 activity in each cell. DMSO: $n$ = 1,724 (iso), 1,187 (hyper); BTZ: $n$ = 1,316 (iso), 1,074 (hyper). Representative data from three independent experiments. The white scale bar represents 200 µm.

(B) Effects of PSMA1 depletion on endogenous ASK activity. The bottom graph depicts the quantification of western blots. Individual values and the mean ± SEM are presented as points and bars, respectively. $n$ = 3.

(C) Effects of proteasome inhibitors on proteasome activity. Cells were treated with 100 nM bortezomib or 10 µM MG132 for indicated time. Mean + SEM. $n$ = 3.

(A, B) Iso, 300 mOsm; Hyper, 400 mOsm; 10 min. $^{***}p < 0.001$.

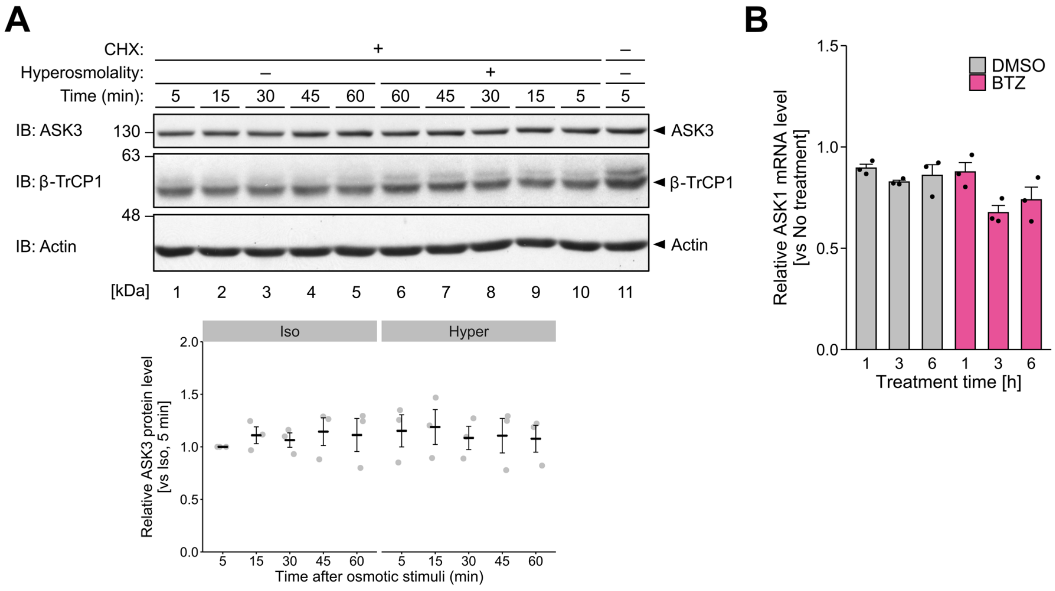


**Figure S3. Involvement of ASK1 protein levels in the regulation of ASK3 signaling under hyperosmotic stress. Related to Figure 3.**

(A) Effects of hyperosmotic stress on the protein level of ASK3 under protein synthesis inhibition. Cells were pretreated with 100 μg/mL cycloheximide for 2 h before osmotic stimuli. The bottom graph depicts the quantification of western blots. Individual values and the mean ± SEM are presented as points and bars, respectively. $n = 3$. Efficiency of protein synthesis inhibition was confirmed by the reduction of β-TrCP1 level. Iso, culture medium; Hyper: culture medium supplemented with 200 mM mannitol. CHX, cycloheximide; IB, immunoblotting.

(B) Effects of proteasome inhibition on ASK1 mRNA levels, related to Figure 3B. Cells were treated with 100 nM bortezomib for the indicated time. Mean + SEM. $n = 3$.

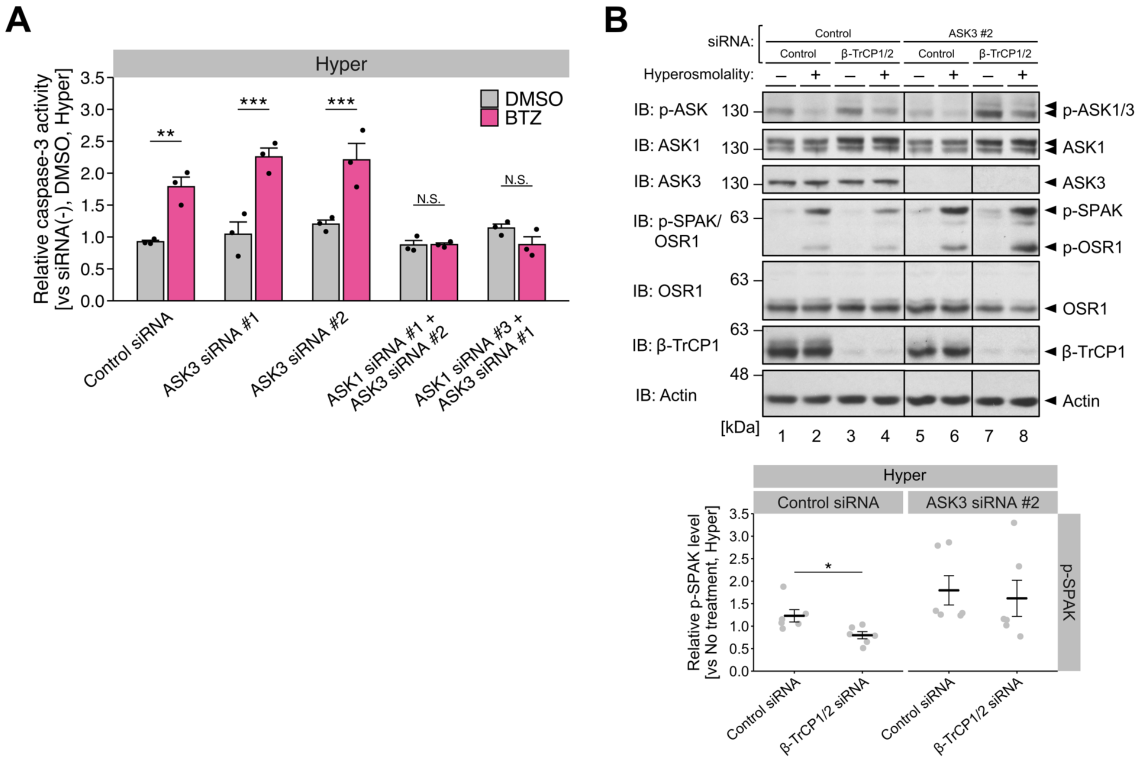


**Figure S4. Involvement of ASK3 in the hyperosmotic stress-induced apoptosis and SPAK/OSR1 activation under ASK1 accumulation.**

(A) Effects of ASK3 and ASK1 depletion on bortezomib-induced overactivation of caspase-3 under hyperosmotic stress, related to Figure 4E. Mean + SEM. $n = 3$. Hyper, culture medium supplemented with 500 mM mannitol; 12 h.

(B) Requirement of ASK3 for the effects of β-TrCP1/2 depletion on SPAK/OSR1 activation under hyperosmotic stress. The bottom graph depicts the quantification of western blots. Individual values and the mean ± SEM are presented as points and bars, respectively. $n = 6$. Iso, 300 mOsm; Hyper, 400 mOsm; 10 min. IB, immunoblotting.

N.S., not significant; $^{*}p < 0.05$, $^{**}p < 0.01$, $^{***}p < 0.001$.

**Table S1. Reagents and Resources used in this study.**

| REAGENT or RESOURCE | SOURCE | IDENTIFIER |
|---|---|---|
| Antibodies | | |
| Mouse monoclonal anti-DYKDDDDK tag (Flag; clone 1E6, IB: 1:5,000–1:10,000) | Wako Pure Chemical Industries | Cat#012-22384; RRID: AB_10659717 |
| Mouse monoclonal anti-Actin (Actin; clone AC-40, IB: 1:2,000) | Sigma-Aldrich | Cat#A3853; RRID: AB_262137 |
| Alexa Fluor 488 goat anti-rat IgG (H + L) (IF: 1:200) | Molecular Probes | Cat#A11006; RRID: AB_2534074 |
| Alexa Fluor 594 goat anti-mouse IgG (H + L) (IF: 1:1,000) | Molecular Probes | Cat#A11005; RRID: AB_2534073 |
| Rabbit monoclonal anti-ASK1 (ASK1; clone EP553Y, IB: 1:5,000–1:10,000) | Abcam | Cat#ab45178; RRID: AB_722915 |
| Rat monoclonal anti-ASK3 (ASK3; IB: 1:5,000–1:10,000) | Naguro et al., 2012[2] | N/A |
| Mouse monoclonal anti-beta-catenin (β-catenin; clone E-5, IB: 1:2,000) | Santa Cruz Biotechnology | Cat#sc-7963; RRID: AB_626807 |
| Rabbit monoclonal anti-β-TrCP (β-TrCP1; clone D13F10, IB: 1:5,000) | Cell Signaling Technology | Cat#4394; RRID: AB_10545763 |
| Mouse monoclonal anti-DYKDDDDK tag (Total-ASK3; clone M2, IF: 1:2,000) | Sigma-Aldrich | Cat#F3165; RRID: AB_259529 |
| Mouse monoclonal anti-OXSR1 (OSR1; clone 2A2-1A2, IB: 1:20,000) | Abnova | Cat#H00009943-M01; RRID: AB_425821 |
| Rabbit polyclonal anti-phospho-ASK (p-ASK1/3; Thr808 in human ASK3 and Thr838 in human ASK1, IB: 1:1,000–1:10,000) | Naguro et al., 2012[2]; Tobiume et al., 2002[3] | N/A |
| Rabbit polyclonal anti-phospho-SPAK/OSR1 (p-SPAK/OSR1; Thr231 in human SPAK and Thr185 in human OSR1, IB: 1:1,000) | Naguro et al., 2012[2] | N/A |
| Rat monoclonal anti-phospho-ASK (Phospho-ASK3; Thr808 in human ASK3, clone PA41, IF: 1:500) | Naguro et al., 2012[2] | N/A |
| Rabbit polyclonal anti-PSMC3 (PSMC3; IB, 1:5,000) | Hamazaki et al., 2007[4] | N/A |
| Mouse monoclonal anti-Ubiquitin (Ubiquitinated proteins; clone P4D1, IB: 1:10,000) | Santa Cruz Biotechnology | Cat#sc-8017; RRID: AB_628423 |
| Anti-rabbit IgG, HRP-linked (IB: 1:1,000–1:20,000) | Cell Signaling Technology | Cat. #7074; RRID: AB_2099233 |
| Anti-rat IgG, HRP-linked (IB: 1:5,000–1:10,000) | Cell Signaling Technology | Cat. #7077; RRID: AB_10694715 |
| Anti-mouse IgG, HRP-linked (IB: 1:2,000–1:20,000) | Cell Signaling Technology | Cat. #7076; RRID: AB_330924 |
| Chemicals, peptides, and recombinant proteins | | |
| 2 × Reaction Buffer | BioVision | Cat#1068 |
| Bortezomib | LC Laboratories | Cat#B-1408 |
| Ac-DEVD-AFC | Cayman | Cat#14459 |
| Dimethyl sulfoxide | Sigma-Aldrich | Cat#D5879 |
| Dithiothreitol | TCI | Cat#D1071 |
| Formaldehyde Solution | Wako Pure Chemical Industries | Cat#064-00406 |
| Hoechst33258 | Dojindo | Cat#343-07961 |
| Phosphatase inhibitor cocktail | Watanabe et al., 2018[5] | N/A |
| MG132 | Enzo Life Sciences | Cat#BML-PI102 |

| | | |
|---|---|---|
| Opti-MEM | Thermo Fisher Scientific | Cat#31985 |
| Polyethylenimine “MAX” | Polysciences | Cat#24765 |
| Adenosine 5'-Triphosphate Disodium Salt Trihydrate | Wako Pure Chemical Industries | Cat#018-16911 |
| Succinyl-Leu-Leu-Val-Tyr-7-amino-4-methylcoumarin | Peptide Institute | Cat#3120-v |
| Lipofectamine RNAiMAX | Invitrogen | Cat#133778-150 |
| Tetracycline | Sigma-Aldrich | Cat#T7660 |
| Dulbecco’s Modified Eagle’s Medium - high glucose | Sigma-Aldrich | Cat#D5796 |
| penicillin G | Meiji Seika | Cat#6111400D2039 |
| Cycloheximide | Sigma-Aldrich | Cat#C7698 |
| Experimental models: Cell lines | | |
| Human: HEK293A cells | Invitrogen | N/A |
| Human: Tetracycline-inducible Flag-ASK3-stably-expressing HEK293A cells | Watanabe et al., 2018[5] | N/A |
| Oligonucleotides | | |
| Control siRNA (Stealth RNAi Negative Control Medium GC Duplex #2) | Invitrogen | Cat#12935-112 |
| ASK1 siRNA #1 (Stealth RNAi siRNA, target sequence: 5'-GCCAACACUACAGUCAGGAAUUAAU-3') | Invitrogen | Cat#10620312 |
| ASK1 siRNA #2 (Stealth RNAi siRNA, target sequence: 5'-CCUGUGCUAACGACUUGCUUGUUGA-3') | Invitrogen | Cat#10620312 |
| ASK1 siRNA #3 (Stealth RNAi siRNA, target sequence: 5'-UGAAGCUAAGUAGUCUUCUUGGUAA-3') | Invitrogen | Cat#10620312 |
| ASK3 siRNA #1 (Stealth RNAi siRNA, target sequence: 5'-CACCGAAGAGCAGUGCAGUAGAUUU-3') | Invitrogen | Cat#10620312 |
| ASK3 siRNA #2 (Stealth RNAi siRNA, target sequence: 5'-GAGAGGGUUUCUUAAGGCAGGUGAA-3') | Invitrogen | Cat#10620312 |
| b-TrCP1 siRNA (Stealth RNAi siRNA, BTRC-HSS113250, target sequence: 5'-CCAACAUGGGCACAUAAACUCGUAU-3') | Invitrogen | Cat#1299001 |
| b-TrCP2 siRNA (Stealth RNAi siRNA, FBXW11-HSS177307, target sequence: 5'-CCAGCCUGGAAUGUUUGAAAGUGUU-3') | Invitrogen | Cat#1299001 |
| Control siRNA (siGENOME Non-targeting siRNA #4, target sequence: 5'-AUGAACGUGAAUUGCUCAA-3') | Horizon | Cat#D-001210-04 |
| PSMA1 siRNA #1 (siGENOME siRNA, target sequence: 5'-GAUAUGGGCCCUCACAUUU-3') | Horizon | Cat#D-010123-01 |
| PSMA1 siRNA #2 (siGENOME siRNA, target sequence: 5'-GGGCAGGAUUCAUCAAAUU-3') | Horizon | Cat#D-010123-03 |
| PSMC2 siRNA #1 (Stealth RNAi siRNA, PSMC2-HSS108700) | Invitrogen | Cat#1299001 |
| PSMC2 siRNA #2 (Stealth RNAi siRNA, PSMC2-HSS108701) | Invitrogen | Cat#1299001 |
| PSMC2 siRNA #3 (Stealth RNAi siRNA, PSMC2-HSS108702) | Invitrogen | Cat#1299001 |
| PSMC3 siRNA #1 (Stealth RNAi siRNA, PSMC3-HSS108703) | Invitrogen | Cat#1299001 |
| PSMC3 siRNA #2 (Stealth RNAi siRNA, PSMC3-HSS108705) | Invitrogen | Cat#1299001 |
| See Table S2 for primers of qPCR analysis. | | |
| Recombinant DNA | | |
| pcDNA4/TO EGFP-Flag-ASK3 KM | Watanabe et al., 2021[6] | N/A |

| | | |
|---|---|---|
| pcDNA3/GW Flag-ASK1 | Takeda et al., 2007[7] | N/A |
| pcDNA3/GW Flag-ASK1 KM | Takeda et al., 2007[7] | N/A |
| pcDNA3/GW Flag-ASK3 | Naguro et al., 2012[2] | N/A |
| pcDNA3/GW EGFP-Flag-ASK1 | This paper | N/A |
| Software and algorithms | | |
| GNU Image Manipulation Program (GIMP; ver. 2.8.22) | GIMP Development Team | https://www.gimp.org/ |
| HCS Studio (ver. 6.4.3) or Cellomics vHCS: View (ver. 1.6.2) | Thermo Fisher Scientific | N/A |
| ImageJ (ver. 1.53k) | Schneider et al., 2012[8] | https://imagej.nih.gov/ij/ |
| Metascape (ver. 3.5) | Zhou et al., 2019[9] | https://metascape.org/ |
| GraphPad Prism 7.0c | GraphPad Software | https://www.graphpad.com/ |
| R (ver. 4.2.2) | R Foundation | https://www.r-project.org/ |
| RStudio (ver. 2022.07.2+576) | RStudio | https://www.rstudio.com/ |

**Table S2. List of the primer sequences used in qPCR analysis.**

| Gene | Forward | Reverse |
|---|---|---|
| human ASK1 | 5'-CACGTGATGACTTAAAATGCTTG-3' | 5'-AGTCAATGATAGCCTTCCACAGT-3' |
| human GAPDH | 5'-AGCCACATCGCTCAGACAC-3' | 5'-GCCCAATACGACCAAATCC-3' |

**Reference**

1. Bader, G.D., and Hogue, C.W.V. (2003). An automated method for finding molecular complexes in large protein interaction networks. BMC Bioinformatics *4*, 2. 10.1186/1471-2105-4-2.

2. Naguro, I., Umeda, T., Kobayashi, Y., Maruyama, J., Hattori, K., Shimizu, Y., Kataoka, K., Kim-Mitsuyama, S., Uchida, S., Vandewalle, A., et al. (2012). ASK3 responds to osmotic stress and regulates blood pressure by suppressing WNK1-SPAK/OSR1 signaling in the kidney. Nat. Commun. *3*, 1285. 10.1038/ncomms2283.

3. Tobiume, K., Saitoh, M., and Ichijo, H. (2002). Activation of apoptosis signal-regulating kinase 1 by the stress-induced activating phosphorylation of pre-formed oligomer. Journal of Cellular Physiology *191*, 95–104. 10.1002/jcp.10080.

4. Hamazaki, J., Sasaki, K., Kawahara, H., Hisanaga, S.-I., Tanaka, K., and Murata, S. (2007). Rpn10-mediated degradation of ubiquitinated proteins is essential for mouse development. Mol. Cell. Biol. *27*, 6629–6638. 10.1128/MCB.00509-07.

5. Watanabe, K., Umeda, T., Niwa, K., Naguro, I., and Ichijo, H. (2018). A PP6-ASK3 Module Coordinates the Bidirectional Cell Volume Regulation under Osmotic Stress. Cell Rep. *22*, 2809–2817. 10.1016/j.celrep.2018.02.045.

6. Watanabe, K., Morishita, K., Zhou, X., Shiizaki, S., Uchiyama, Y., Koike, M., Naguro, I., and Ichijo, H. (2021). Cells recognize osmotic stress through liquid-liquid phase separation lubricated with poly(ADP-ribose). Nat. Commun. *12*, 1353. 10.1038/s41467-021-21614-5.

7. Takeda, K., Shimozono, R., Noguchi, T., Umeda, T., Morimoto, Y., Naguro, I., Tobiume, K., Saitoh, M., Matsuzawa, A., and Ichijo, H. (2007). Apoptosis signal-regulating kinase (ASK) 2 functions as a mitogen-activated protein kinase kinase kinase in a heteromeric complex with ASK1. J. Biol. Chem. *282*, 7522–7531. 10.1074/jbc.M607177200.

8. Schneider, C.A., Rasband, W.S., and Eliceiri, K.W. (2012). NIH Image to ImageJ: 25 years of image analysis. Nat. Methods *9*, 671–675. 10.1038/nmeth.2089.

9. Zhou, Y., Zhou, B., Pache, L., Chang, M., Khodabakhshi, A.H., Tanaseichuk, O., Benner, C., and Chanda, S.K. (2019). Metascape provides a biologist-oriented resource for the analysis of systems-level datasets. Nat. Commun. *10*, 1523. 10.1038/s41467-019-09234-6.
